# A coupled cerebro-ocular-CSF lumped-parameter model under gravitational and postural variations

**DOI:** 10.64898/2026.03.17.712384

**Authors:** Michele Nigro, Andrea Montanino, Eduardo Soudah

## Abstract

Spaceflight-Associated Neuro-ocular Syndrome (SANS) involves complex interactions between intracranial pressure (ICP), intraocular pressure (IOP), and cerebrospinal fluid (CSF) dynamics within the optic nerve subarachnoid space (ONSAS). While existing computational models address specific aspects of these interactions, they lack a comprehensive, system-level representation. To bridge this gap, we present the HEAD (Hemodynamic Eye-brain Associated Dynamics) model. By consistently integrating several previously proposed physiological sub-models, HEAD provides a unified lumped-parameter framework that fully couples cerebrovascular autoregulation, multi-territory ocular hemodynamics, and compartmentalized craniospinal-ONSAS CSF circulation under gravitational loading. This formulation enables the simultaneous analysis of eye–brain–CSF dynamics within a single computational tool. Model predictions were validated against experimental data from supine (0°) to head-down tilt (HDT, −30°) postures, accurately reproducing posture-dependent IOP increases and achieving an excellent ICP match against clinical benchmarks at the −6° HDT standard bed-rest angle. The coupled system predicts bed-specific ocular hemodynamic responses, with retinal blood flow exhibiting the largest relative increase under HDT compared to the ciliary and choroidal circulations. Crucially, explicitly modeling the ONSAS as a distinct compartment reveals a posture-dependent pressure drop of 1.89–3.69 mmHg between the intracranial and perioptic spaces. This compartmentalization yields a translaminar pressure profile that remains positive (8.05–11.83 mmHg) across all simulated conditions but is chronically reduced under sustained HDT. Ultimately, the HEAD model elucidates the physiological mechanisms linking gravitational stress to translaminar mechanics, providing a robust computational foundation to investigate SANS and supply boundary conditions for structural models of the optic nerve head.

## 1 Introduction

Spaceflight-Associated Neuro-ocular Syndrome (SANS) has emerged as one of the most significant physiological risks associated with long-duration human spaceflight. Astronauts exposed to microgravity frequently develop neuro-ophthalmic alterations including optic disc edema, posterior globe flattening, choroidal folds, hyperopic refractive shifts, and optic nerve sheath distension [1, 2]. The incidence of early optic disc edema in long-duration missions reaches up to ∼70%, highlighting the need for improved mechanistic understanding and effective mitigation strategies [3, 4]. Although the precise mechanisms underlying SANS remain incompletely understood, current evidence suggests that the syndrome arises from complex interactions between vascular, cerebrospinal fluid (CSF), and ocular pressure dynamics that are profoundly altered in the microgravity environment.

One of the primary physiological changes observed during spaceflight is the redistribution of body fluids toward the head due to the absence of hydrostatic pressure gradients. This headward fluid shift is believed to affect intracranial pressure (ICP), ocular venous drainage, and the pressure balance across the lamina cribrosa [5, 2]. As a consequence, the pressure environment surrounding the optic nerve head may be substantially modified. In particular, the mechanical balance across the lamina cribrosa, commonly described through the translaminar pressure difference between intraocular pressure (IOP) and the retrolaminar CSF pressure, plays a critical role in determining the mechanical loading of the optic nerve head and the perfusion of the ocular circulation. Changes in this balance have been hypothesized to contribute to the development of optic disc edema and other structural alterations observed in astronauts.

Recent experimental and imaging studies have suggested that the CSF contained within the optic nerve subarachnoid space (ONSAS) may behave as a partially compartmentalized system rather than a simple extension of the intracranial CSF space. In particular, posture-dependent compartmentalization of CSF within the ONSAS has emerged as a potential contributor to SANS-related structural changes [5, 6]. Such compartmentalization may arise from the complex geometry of the optic nerve sheath and from mechanical interactions between the lamina cribrosa, the surrounding tissues, and the perioptic CSF space. Under certain conditions, these factors may allow the retrolaminar CSF pressure to deviate from intracranial pressure, thereby modifying the translaminar pressure balance. Beyond spaceflight, the interaction between IOP and retrolaminar CSF pressure has also been implicated in terrestrial pathologies such as normal-tension glaucoma [7, 8] and idiopathic intracranial hypertension [9]. These observations highlight the importance of the coupled eye–brain pressure environment for both space medicine and clinical ophthalmology.

Because direct measurements of these pressure interactions are challenging in vivo, computational modeling has become an important tool for investigating the underlying mechanisms of SANS. Experimental studies rely on in-flight measurements and ground-based analogs such as head-down tilt (HDT) and dry immersion, which reproduce some aspects of cephalad fluid shifts but cannot fully replicate the long-term physiological adaptations associated with microgravity [10, 11, 12, 13]. Mathematical models provide a complementary framework in which the complex interactions between vascular dynamics, ocular physiology, and CSF circulation can be analyzed under controlled conditions [14, 15, 16]. Despite these advances, most computational approaches address only subsets of the coupled eye–brain dynamics. Some models reproduce posture-dependent intraocular pressure variations but neglect cerebrovascular regulation or CSF compartmentalization [15, 17]. Other models incorporate systemic cardiovascular interactions without explicitly resolving ocular hemodynamics or ONSAS dynamics. The eye–brain model proposed by Salerni et al. [18] represents three ocular vascular beds coupled to a simplified brain module, but assumes that the intracranial pressure is identical to the CSF pressure in the optic nerve sheath (*P*_ONSAS_) and does not resolve the detailed cerebrovascular circulation. Multiscale cardiovascular frameworks including cerebral autoregulation and globe-level IOP modules [19, 14, 20] capture closed-loop hemodynamics but do not resolve bed-specific ocular blood flows or perioptic CSF dynamics. Conversely, the CSF compartmentalization model proposed by Holmlund et al. [21] describes posture-dependent ONSAS resistance through Darcy flow in the optic nerve sheath, yet does not include ocular hemodynamics or cerebrovascular autoregulation. As a result, currently available lumped or multiscale models are unable to self-consistently compute the bidirectional coupling between intraocular pressure and ONSAS pressure under varying gravitational conditions.

In this work, we hypothesize that representing the ONSAS as a posture-sensitive CSF compartment dynamically distinct from the intracranial CSF space alters the predicted retrolaminar pressure environment and therefore modifies both the translaminar pressure balance and the ocular hemodynamic response during gravitational transitions. To investigate this hypothesis, we introduce the HEAD (Hemodynamic Eye-brain Associated Dynamics) model, a unified lumped-parameter framework that integrates cerebrovascular circulation, ocular hemodynamics, and craniospinal CSF dynamics into a single coupled system of ordinary differential equations. This model combines established physiological formulations while introducing a set of structural extensions that allow the explicit representation of eye–brain pressure interactions. In particular, the framework incorporates cerebrovascular dynamics with autoregulation and a resolved Circle of Willis, multi-territory ocular hemodynamics, including retinal, choroidal, and ciliary vascular beds coupled with lamina cribrosa mechanics, and a compartmentalized representation of craniospinal and perioptic CSF circulation in which the ONSAS can dynamically decouple from intracranial pressure.

The objectives of the present study are threefold. First, we validate the coupled model against experimental measurements of intraocular and intracranial pressure across four tilt conditions spanning from supine 0° to −30° head-down tilt. Second, we characterize the bed-specific ocular hemodynamic response of the retinal, choroidal, and ciliary circulations under progressive head-down tilt, a prediction that cannot be obtained from models that treat the ocular circulation as a single compartment. Third, we quantify posture-dependent ONSAS compartmentalization and its effect on the translaminar pressure balance and on CSF exchange flows through the optic nerve head. By providing a physiologically consistent description of the coupled eye–brain–CSF dynamics under varying gravitational conditions, the proposed framework aims to contribute to a better mechanistic understanding of the pressure environment associated with SANS and related neuro-ophthalmic alterations.

This paper is organized as follows. Section 2 presents the model formulation and coupling strategy. Section 3 reports the simulation results and model validation. Section 4 and Section 5 discuss the physiological implications of the findings, as well as the limitations of the present framework and directions for future work. For completeness, the full set of governing equations and parameter values for the cerebrovascular subsystem is provided in the Appendix.

## 2 Materials and Methods

The circulatory and fluid-mechanical systems of the eye and brain are strongly coupled: they share arterial supply through the ophthalmic artery and mechanical loading through the CSF pressure transmitted to the ONSAS. Changes in body orientation modify the hydrostatic pressure gradients acting on these interconnected compartments, making an integrated modeling approach essential to capture their coupled responses.

The HEAD model is formulated as a lumped-parameter (0D) representation of the coupled cerebrovascular, ocular, and CSF systems. In this framework, spatially distributed vascular and CSF networks are represented as interconnected compartments governed by ODEs. Each compartment is described by a set of variables including pressure, volume, and flow. The model equations follow the classical cardiovascular-electrical analogy, in which pressure corresponds to voltage, flow rate to electrical current, viscous resistance to electrical resistance, and vascular or CSF compliance to capacitance [22, 23, 24]. This formulation, widely used in cerebrovascular and ocular modeling [25, 26, 27], enables rapid parametric studies, direct comparison with lumped experimental observables (IOP, ICP, blood flow), and straightforward implementation of gravitational offsets through hydrostatic pressure terms.

The model comprises three coupled systems: (i) a cerebrovascular system, with a model of the brain circulation, including the Circle of Willis and cerebral autoregulation mechanisms [25]; (ii) an ocular hemodynamic system, including the complete ocular blood circulation, the lamina cribrosa dynamics, the aqueous liquid, and ocular volumes dynamics [28, 17, 18], and (iii) a craniospinal and ONSAS CSF dynamics system. These are coupled through shared boundary pressures at physiologically defined interfaces, described in Section 2.4. The individual subsystems are described in the following sections, while the complete mathematical formulation is detailed in the appendix. The complete model schematic is shown in Fig.1, Fig.2, and Fig.3.

**Figure 1:**
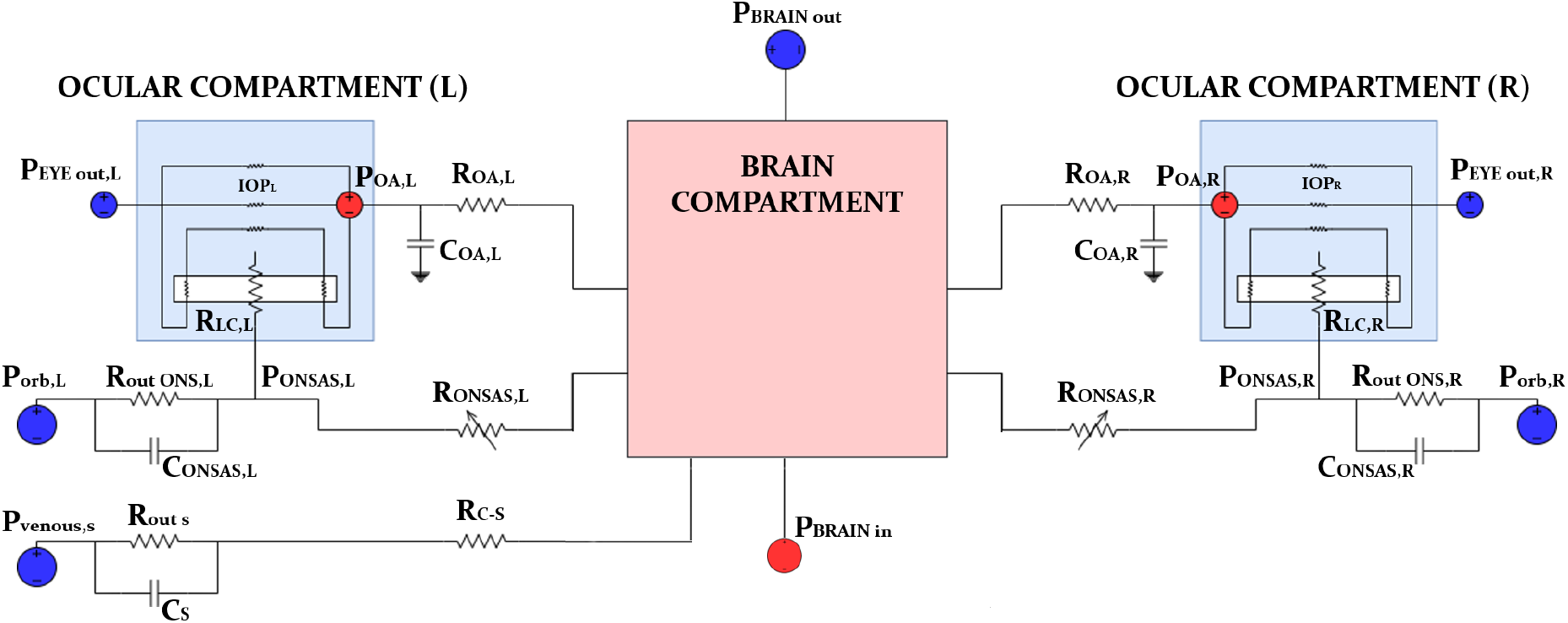
Coupled eye-brain-CSF model architecture. Lumped-parameter schematic of the integrated model, comprising a central brain compartment coupled to left and right ocular compartments and to the optic-nerve subarachnoid space (ONSAS) dynamics. The ophthalmic arterial pressure *P*_*OA*_ provides the hemodynamic input to each eye through an ophthalmic artery resistance-compliance element (*R*_*OA*_, *C*_*OA*_). Each ocular module generates the intraocular pressure (IOP) and exchanges stresses and pressures with the local ONSAS pressure *P*_*ONSAS*_ at the lamina cribrosa via an effective laminar element *R*_*LC*_. The ONSAS compartment is modeled as a distinct cerebrospinal fluid (CSF) space with its own compliance *C*_*ONSAS*_ and outflow pathway *R*_*o*ut,ONS_, referenced to an orbital or venous pressure *P*_*orb*_, allowing *P*_*ONSAS*_ to differ from intracranial pressure and to vary with posture. Red and blue boundary nodes denote imposed arterial inflow and venous or outflow reference pressures, respectively; resistive, compliant, and variable elements represent viscous losses, compartmental compliances, and pressure-dependent collapsible segments.

**Figure 2:**
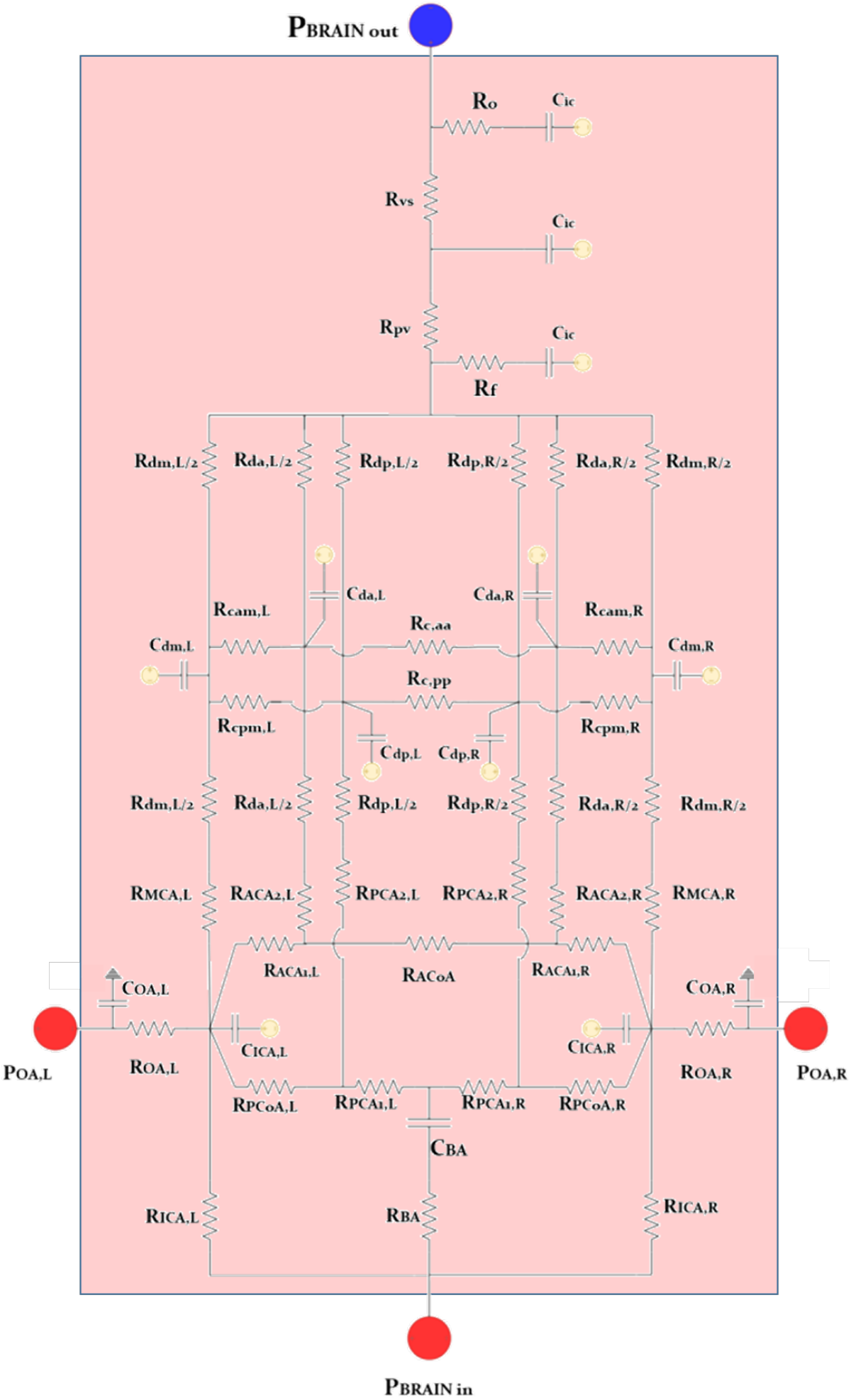
Cerebrovascular subsystem: Circle of Willis, distal territories, and intracranial-CSF coupling. Circuit representation of the cerebral arterial network, including bilateral internal carotid arteries (ICA), basilar artery (BA), and Circle of Willis pathways (ACA, MCA, and PCA branches with communicating arteries), followed by distal vascular territories modeled through compartmental resistances and compliances. The model resolves left-right flow redistribution through the communicating segments and incorporates distal pressure-flow dynamics via lumped arterial and downstream elements. The intracranial cerebrospinal fluid (CSF) module represents CSF formation, storage, and outflow pathways through resistive elements and intracranial compliance, providing a dynamic coupling between intracranial pressure and cerebral venous drainage. Red boundary nodes denote imposed arterial inputs, whereas the blue node denotes the venous or sinus reference pressure; capacitors represent vascular and compartmental compliance, and resistors represent viscous losses.

**Figure 3:**
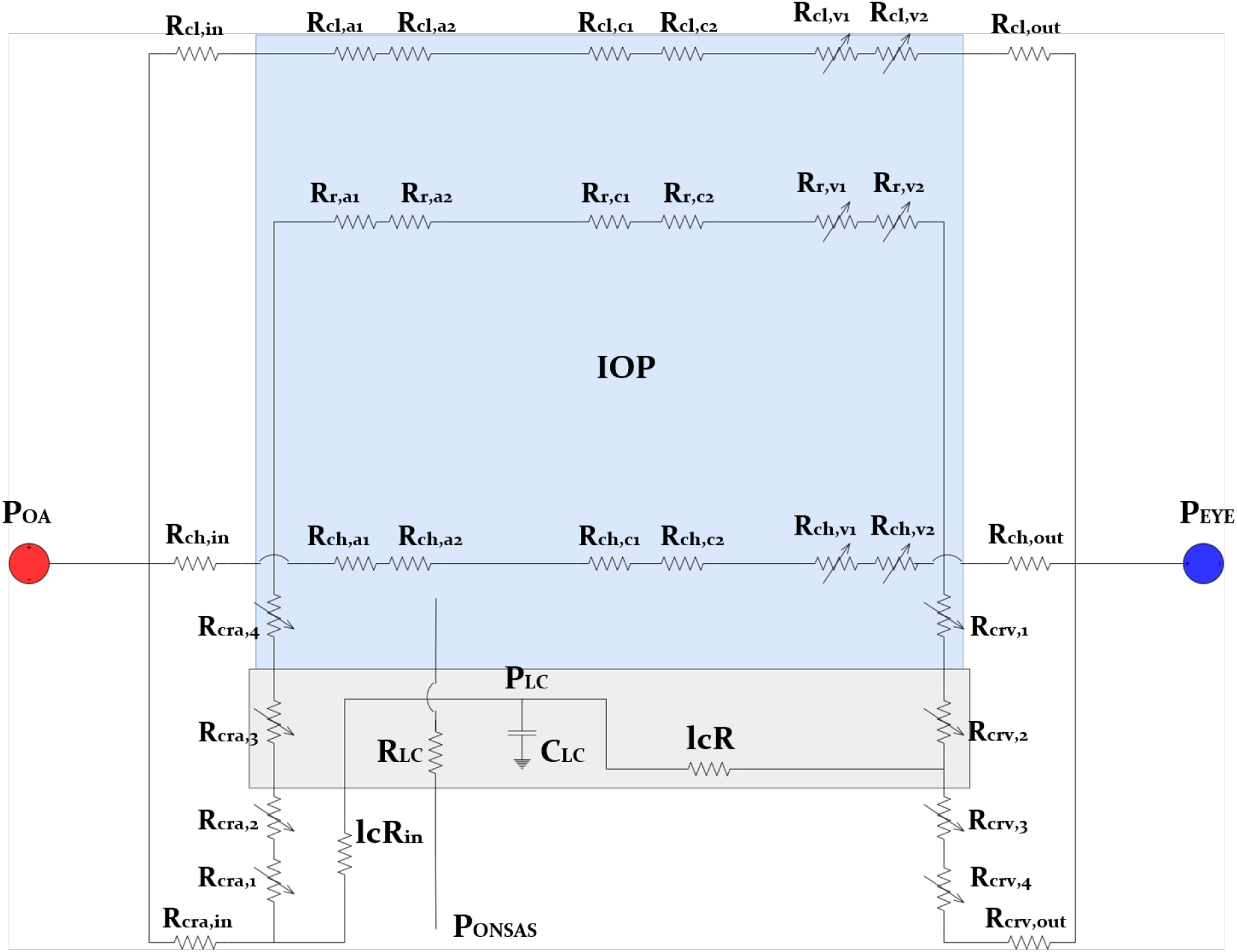
Ocular subsystem: retinal and choroidal circulations coupled to intraocular and ONSAS pressures. Detailed electrical-analogy circuit of the ocular hemodynamic module. The ophthalmic arterial pressure *P*_*OA*_ drives blood flow through the central retinal artery (CRA) and through the microvascular compartments of the retina (*r, a*; *r, c*; *r, v*) and choroid (*ch, a*; *ch, c*; *ch, v*), with venous drainage referenced to an episcleral or venous outlet pressure *P*_*EYE*_. Intraocular pressure (IOP) acts as an external pressure on intraocular vascular segments, while the optic-nerve subarachnoid space pressure *P*_*ONSAS*_ acts on retrolaminar (retrobulbar) segments, enabling posture-dependent modulation of transmural pressure. The lamina cribrosa is represented as an intermediate compartment with pressure *P*_*LC*_ that mediates eye-ONSAS coupling and defines the translaminar mechanical loading. Variable resistances indicate passive pressure-dependent changes in vascular caliber induced by transmural pressure variations.

### 2.1 Cerebrovascular subsystem

The cerebrovascular system builds upon the multicomponent model of Ursino and Giannessi [25], which provides a lumped-parameter (0D) representation of the large cerebral arteries, the Circle of Willis (CoW), and the intracranial CSF-venous system. The arterial network includes the bilateral internal carotid arteries (ICA), the basilar artery (BA), and the six major branches of the Circle of Willis: the anterior cerebral arteries (ACA), middle cerebral arteries (MCA), and posterior cerebral arteries (PCA), interconnected through the anterior and posterior communicating arteries (ACoA and PCoA). Each large artery is modeled as a resistive-compliant element whose baseline resistance is estimated from the Hagen-Poiseuille law using anatomical lengths and radii reported in the literature. Distal to the CoW, the parenchymal microcirculation is represented through pressure-dependent compartmental resistances and compliances. Cortical anastomoses connect the anterior, middle, and posterior territories bilaterally, enabling the model to reproduce collateral flow redistribution during changes in cerebral perfusion pressure [25]. Cerebral autoregulation is implemented through two coupled control mechanisms: a myogenic response proportional to deviations of cerebral perfusion pressure from its normotensive set point, and a metabolic reactivity mechanism driven by arterial CO_2_ tension. Both mechanisms act by modifying the effective radius of distal arteriolar compartments, thereby altering vascular resistance according to Poiseuille’s law. The intracranial compartment is coupled to a lumped CSF balance including production, absorption toward the venous sinuses, and an intracranial compliance *C*_*ic*_ relating ICP dynamics to variations in intracranial blood volume.

In the HEAD model, the original Ursino-Giannessi architecture is extended in two key ways to enable integrated eye-brain-CSF simulations under gravitational stress. First, the cerebrovascular model is expanded to explicitly represent the bilateral vascular territories of the Circle of Willis, introducing a territory index *j* ∈ MCA, ACA, PCA and a hemispheric index *s* ∈ *L, R* to account for lateralized cerebral perfusion. Second, two structural modifications are introduced to couple the cerebrovascular system with the ocular and CSF subsystems. With the objective of enhancing ICP dynamics, the governing equation for intracranial pressure is augmented by additional CSF exchange terms derived from the craniospinal-ONSAS module. In particular, the intracranial volume balance includes the cranio-spinal flow *Q*_*c*-*s*_ and the bilateral cranio-ONSAS flows *Q*_*c*-ONSAS,*L*_ and *Q*_*c*-ONSAS,*R*_. These terms extend the original Ursino formulation to capture posture-dependent CSF redistribution, a key mechanism in SANS-related physiology. Furthermore, the mass balance equations of the internal carotid arteries are augmented by an ophthalmic extraction flow

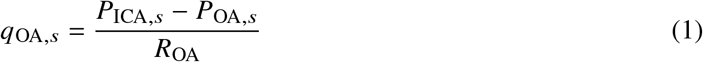

This term represents blood diversion toward the ophthalmic artery and establishes a direct hemodynamic coupling between cerebral and ocular circulation. Such ocular-cerebral hemodynamic coupling is absent in the original Ursino-Giannessi formulation, which does not include a dedicated ophthalmic outflow pathway.

Together, these architectural extensions enable the HEAD model to capture the bidirectional coupling between cerebral circulation, ocular hemodynamics, and CSF dynamics, providing a unified platform to investigate eye-brain-CSF interactions under posture-dependent gravitational conditions.

### 2.2 Ocular hemodynamic subsystem

While the cerebrovascular subsystem governs cerebral circulation and intracranial pressure regulation, the ocular subsystem resolves the vascular, mechanical, and fluid dynamics within the eye and its interaction with the optic nerve head. The ocular hemodynamic subsystem includes the full ocular vascular circulation together with lamina cribrosa mechanics and intraocular pressure dynamics.

#### 2.2.1 Ocular vascular circuit

The ocular hemodynamic subsystem resolves blood flow through three parallel vascular territories (retinal, choroidal, and ciliary), each represented by a series of resistive compartments corresponding to successive vascular segments: arterioles (a), capillaries (c), and two venular stages (v_1_, v_2_). The vascular architecture follows the lumped-parameter framework originally proposed by Guidoboni et al. [26] and later extended by Salerni et al. [18] to incorporate coupling with cerebral circulation. In the HEAD model, this formulation is further extended to establish a fully integrated eye-brain coupling. Blood enters the ocular circulation through the ophthalmic artery, modeled as a resistive-compliant element (*R*_*OA*_, *C*_*OA*_) whose input pressure *P*_*OA*_ is dynamically provided by the cerebrovascular subsystem. Venous drainage from all vascular territories converges to a common outlet pressure *P*_*OUT,EYE*_ representing the ocular venous pressure.

A key feature of the proposed HEAD model is the explicit distinction between intraocular and retrolaminar vessel segments. Vessels located within the globe are subjected to IOP as the external compressive load, whereas vessels located posterior to the lamina cribrosa experience the pressure within the optic nerve subarachnoid space (*P*_*ONSAS*_). This anatomical separation enables the model to capture posture-dependent mechanical loading of the retinal venous outflow pathway.

The corresponding vascular resistances are therefore modeled as passive, transmural-pressure-dependent elements. Variable resistances are applied to venular segments in the retinal and choroidal territories, as well as to the CRA and CRV segments crossing the lamina cribrosa. The resistance law follows a sigmoidal function of transmural pressure, ensuring physiological vessel patency while reproducing the progressive increase in resistance under IOP elevation or retrolaminar compression [26]. This formulation allows the HEAD model to capture flow-limiting venous collapse mechanisms that are particularly relevant under altered gravitational loading.

#### 2.2.2 Lamina cribrosa

The lamina cribrosa is modeled as an intermediate compartment with pressure *P*_*LC*_ located between the retinal vasculature and the ONSAS, following the approach proposed and validated by Sala et al. [28]. This compartment is characterized by a compliance *C*_*LC*_, an inflow resistance *lcR*_*in*_ from the retinal circulation, and an outflow resistance *lcR* connecting it to the ONSAS compartment. Within the HEAD model, this compartment plays a central role in mediating the mechanical interaction between ocular and CSF pressures. In particular, it enables the dynamic computation of the translaminar pressure difference

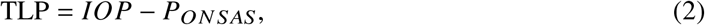

as well as the associated translaminar CSF exchange through the optic nerve head. This coupling provides a mechanistic link between ocular mechanics and CSF dynamics, which is considered relevant for glymphatic transport and for the pathophysiology of SANS-related optic nerve alterations.

#### 2.2.3 Intraocular pressure dynamics

Finally, the IOP is determined by the balance between aqueous humor production and drainage, coupled with the mechanical response of the ocular globe to volume variations. This component follows the globe-level lumped-parameter framework originally developed by Nelson et al. [17] and later adapted by Petersen et al. [15] to investigate posture-dependent ocular dynamics. Within the HEAD model, the IOP module is dynamically coupled to both the ocular vascular circulation and the ONSAS pressure, enabling the consistent computation of intraocular pressure under gravitational reorientation. This coupling allows the model to capture the combined influence of vascular, mechanical, and CSF-related mechanisms on IOP regulation.

### 2.3 Cerebrospinal fluid (CSF) dynamics and ONSAS

The CSF subsystem describes the production, circulation, and absorption of cerebrospinal fluid across the intracranial, spinal, and bilateral ONSAS compartments. Its formulation integrates elements from the Ursino and Giannessi cerebrovascular model [25], which governs CSF production and intracranial pressure dynamics, with the compartmental craniospinal-ONSAS model proposed by Holmlund et al. [21]. This integration enables the consistent representation of CSF redistribution across the craniospinal axis and the optic nerve sheath within the unified HEAD model (Fig. 1).

CSF formation is not prescribed as a constant rate but arises naturally from the cerebrovascular hemodynamics. Specifically, it is modeled as a pressure-dependent filtration from the cerebral capillary bed (pressure *P*_*c*_) into the intracranial compartment (pressure *ICP*) through a formation resistance *R* _*f*_, yielding a production rate

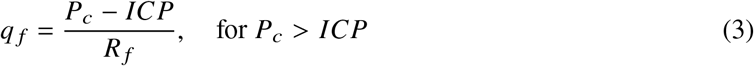

as proposed by Ursino and Giannessi [25]. This formulation ensures that CSF production responds dynamically to changes in cerebral perfusion pressure induced by posture or gravitational loading, in contrast to models that assume a fixed secretion rate. CSF absorption from the cranial compartment toward the dural venous sinuses is driven by the pressure difference between *ICP* and the venous sinus pressure *P*_*vs*_ through an outflow resistance *R*_*o*_ [25]. The intracranial compliance *C*_*ic*_, governing the relationship between intracranial volume changes and ICP, is modeled as a nonlinear function of pressure

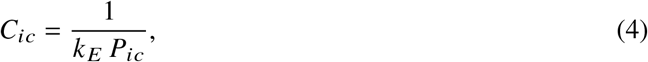

consistent with classical craniospinal compliance observations [29].

From the cranial compartment, CSF is distributed through three parallel pathways: the spinal subarachnoid space, the left ONSAS, and the right ONSAS.

The spinal subarachnoid space is connected to the cranial compartment through a craniospinal resistance *R*_*C*-*S*_, with flow driven by the pressure difference between *ICP* and the spinal CSF pressure *P*_*s*_. The spinal pressure-volume relationship follows a nonlinear exponential elastance formulation with coefficient *E* [21]. CSF is absorbed from the spinal compartment toward the spinal venous pressure *P*_*vs*_ through an outflow resistance *R*_*out,s*_.

While the spinal pathway relies on well-established formulations, a central methodological innovation of the HEAD model lies in its treatment of the perioptic pathways. Rather than assuming instantaneous pressure equilibration between the brain and the eye (i.e., *P*_ONSAS_ = ICP) as in most previous lumped-parameter models [17, 18, 14, 30], we explicitly represent the left and right ONSAS as two distinct compliant compartments.

This distinction is physiologically motivated by experimental evidence showing posture-dependent compartmentalization of CSF within the optic nerve sheath. In particular, upright postures can induce sheath collapse and microstructural obstruction that partially decouple ONSAS pressure from intracranial pressure [21, 7, 31].

Each ONSAS compartment receives CSF from two sources: (i) a cranio-ONSAS flow *Q*_*C*-*ONSAS*_ driven by the pressure difference between *ICP* (corrected for the hydrostatic column *ρ*_*cs f*_ *g h*_*c*-*lc*_ between the cranial reference and the lamina cribrosa) and *P*_*ONSAS*_ through the axial ONSAS resistance *R*_*ONSAS*_; and (ii) a translaminar flow *Q*_*LC*_ driven by the pressure difference between intraocular pressure (IOP) and *P*_*ONSAS*_ through the lamina cribrosa resistance *R*_*LC*_.

CSF exits the ONSAS through trans-sheath absorption toward an orbital reference pressure *P*_*orb*_ through the optic nerve sheath resistance *R*_*out,ONS*_ [21]. The ONSAS resistance *R*_*ONSAS*_ is computed using a Darcy-flow formulation for the porous CSF space within the optic nerve sheath [21]. The sheath is subdivided into four anatomical regions (bulbar, mid-orbital_1_, mid-orbital_2_, and canalicular), and the total resistance is obtained as the sum of regional contributions depending on local sheath geometry, CSF viscosity, porosity, and permeability.

Both the ONSAS resistance *R*_*ONSAS*_ and the ONSAS compliance *C*_*ONSAS*_ are posture-dependent quantities. The local sheath radius varies with *P*_*ONSAS*_ according to MRI-derived distensibility values [21]. Consequently, upright postures induce sheath narrowing, increased hydraulic resistance, and reduced compliance, reproducing the experimentally observed CSF compartmentalization within the optic nerve sheath.

Together, these formulations define the craniospinal-ONSAS CSF subsystem within the HEAD model, whose pressure and flow dynamics are bidirectionally coupled with the cerebrovascular and ocular subsystems through physiologically defined interfaces described in the following section.

### 2.4 Model coupling and numerical implementation

The previous sections describe the physiological formulation of the three subsystems composing the HEAD model. In this section, we present how these components are numerically integrated into a single coupled system, including the definition of the pressure interfaces that connect the cerebrovascular, ocular, and CSF compartments. To couple these three subsystems, we have defined the following physiological interfaces that enforce bidirectional pressure exchange (Fig. 1):

#### Ophthalmic artery

The ocular arterial input pressure *P*_*OA*_ is derived from the cerebral arterial pressure at the level of the internal carotid artery. This connection represents the hemodynamic link between the cerebral circulation and the ocular vascular network.

#### Lamina cribrosa-ONSAS interface

The lamina cribrosa plays a dual role. From the hemodynamic standpoint, the laminar tissue pressure *P*_*LC*_ is governed by a lumped RC equation driven by blood inflow from the retrobulbar arterial input and venous outflow through the post-laminar CRV segment. In this context, *P*_*LC*_ acts as the external compressive pressure on the translaminar portions of the CRA and CRV, thereby modulating their passive resistance. From the CSF standpoint, the lamina acts as a porous membrane permitting a small aqueous-humor outflow *Q*_*LC*_ from the eye into the ONSAS, thereby coupling intraocular and retrobulbar CSF dynamics.

#### ONSAS-intracranial CSF interface

The ONSAS pressure *P*_*ONSAS*_ is dynamically coupled to the ICP through the cranio-ONSAS pathway, allowing pressure and flow interactions between the perioptic CSF space and the intracranial compartment.

These interfaces allow us to define a fully coupled lumped-parameter system composed of 45 nonlinear ordinary differential equations (ODEs) describing the eye-brain-CSF dynamics model. The model was implemented in MATLAB^®^ (R2023a, MathWorks, Natick, MA) and integrated using the stiff solver ode15s, a variable-order implicit method based on backward differentiation formulas and well suited for multicompartment physiological systems with widely separated time scales. Details of the full mathematical equations and geometric parameters used to characterize the systems are provided in Appendix A.

#### 2.4.1 Gravitational simulations

To ensure physical consistency across the model, a unified gravitational framework is introduced in which posture-dependent hydrostatic corrections are applied simultaneously to the hemodynamic boundary conditions and to the CSF intercompartmental flows. Because arterial blood, venous blood, intracranial and spinal CSF, and the ONSAS fluid are all subject to hydrostatic loading, changes in body orientation modify the effective driving pressures throughout the entire system. Gravitational reorientation introduces a hydrostatic pressure correction of the form:

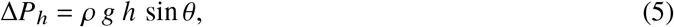

where *ρ* denotes the relevant fluid density (blood or CSF), *g* is the gravitational acceleration, *h* is the vertical anatomical distance between the compartment reference point and a chosen datum, and *θ* is the body tilt angle measured from the horizontal (positive upward, with *θ* < 0° corresponding to HDT).

The primary hemodynamic input to the model is the arterial pressure *P*_in_ prescribed at the level of the brachiocephalic trunk. This location represents the last well-defined upstream node before the cerebral and ocular circulations diverge and allows direct interfacing with whole-body cardiovascular models. The baseline value of *P*_in_ for the supine condition (*θ* = 0°) was obtained from the 0D cardiovascular model of Heldt et al. [32, 33] evaluated at the brachiocephalic compartment.

For each tilt angle *θ*, the arterial input pressure is corrected for the hydrostatic column between the heart and the cranial reference level:

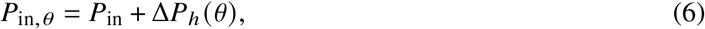

where Δ*P*_*h*_ (*θ*) represents the posture-dependent hydrostatic offset. By convention, head-down tilt (*θ* < 0°) produces a positive offset, reproducing the cephalad pressure increase observed experimentally during HDT maneuvers. To compare model predictions with available experimental analog data, four tilt conditions were simulated: *θ* = 0° (supine), −6°, −15°, and −30° (Fig. 4), spanning the range commonly adopted in ground-based spaceflight analog studies [15, 14].

**Figure 4:**
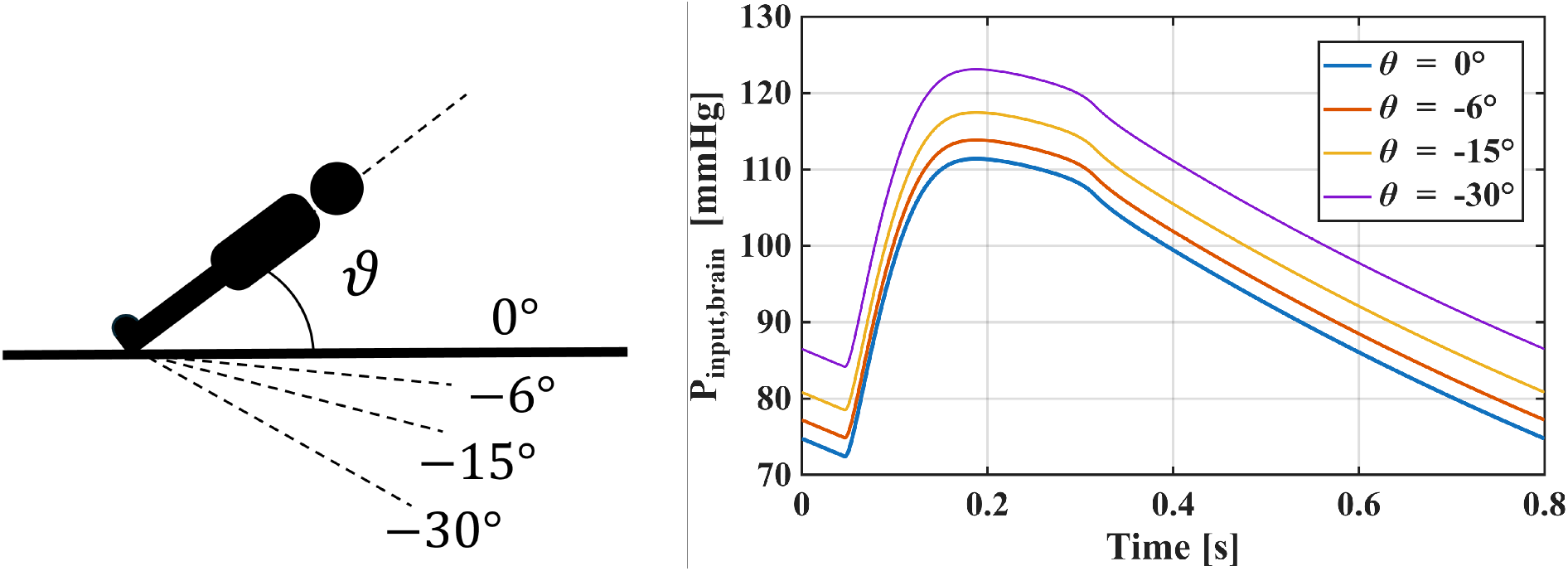
On the left, definition of the tilt angle *θ* (supine *θ* = 0°, head-down tilt *θ* < 0°). On the right, representative pulsatile waveform of the resulting cranial arterial input pressure, *P*_in,*θ*_, at each tilt angle considered, shown over one cardiac cycle and used as upstream forcing for the coupled cerebrovascular and ocular subsystems.

The posture-dependent redistribution of CSF between the intracranial, spinal, and ONSAS compartments follows the approach proposed by Holmlund et al. [21]. In this framework, the hydrostatic column modifies the effective pressure gradients that drive CSF flow between compartments through geometric factors determined by the anatomical distances between reference locations.

The coupled eye-brain-CSF subsystems enables the evaluation of two physiological metrics relevant for the study of SANS:

- **Translaminar Pressure (TLP):** representing the pressure difference across the lamina cribrosa that determines the retrolaminar mechanical loading acting on the optic nerve head.

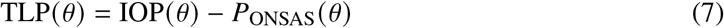
- **Ocular Perfusion Pressure (OPP):** representing the effective upstream pressure available to drive ocular blood flow.

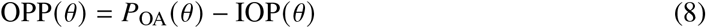

Importantly, these quantities arise directly from the coupled formulation rather than being imposed as boundary conditions. In contrast to previous cerebro-ocular frameworks, the present model explicitly represents the ONSAS as a posture-dependent CSF compartment dynamically distinct from the intracranial space, allowing the retrolaminar pressure entering the TLP definition to be computed as *P*_ONSAS_ rather than assumed equal to ICP. Furthermore, the ophthalmic arterial pressure *P*_OA_ is not prescribed a priori but emerges from the coupling between the cerebral arterial circulation and the ocular vasculature through the resistive-compliant ophthalmic artery element (*R*_*O*A_, *C*_OA_). As a result, the predicted OPP reflects the dynamically transmitted pressure available to drive ocular perfusion under postural and gravitational changes. Taken together, these formulations define the complete HEAD model used to investigate the coupled eye-brain-CSF dynamics under posture-dependent conditions.

## 3 Results

The following section presents the predictions of the HEAD model to quantify the effects of posture-dependent gravitational loading on the coupled eye-brain-CSF dynamics.

### 3.1 Validation of IOP and ICP across postural changes

The construction of the experimental benchmarks and the statistical procedures used for model-data comparison are detailed in Appendix B. Briefly, IOP validation relied on subject-level measurements, whereas ICP benchmarks were assembled by angle-wise pooling of study-level means from the literature. Table 1 summarizes the quantitative comparison, while Fig. 5 reports the corresponding experimental distributions (IOP) and pooled study benchmarks (ICP).

**Table 1:**
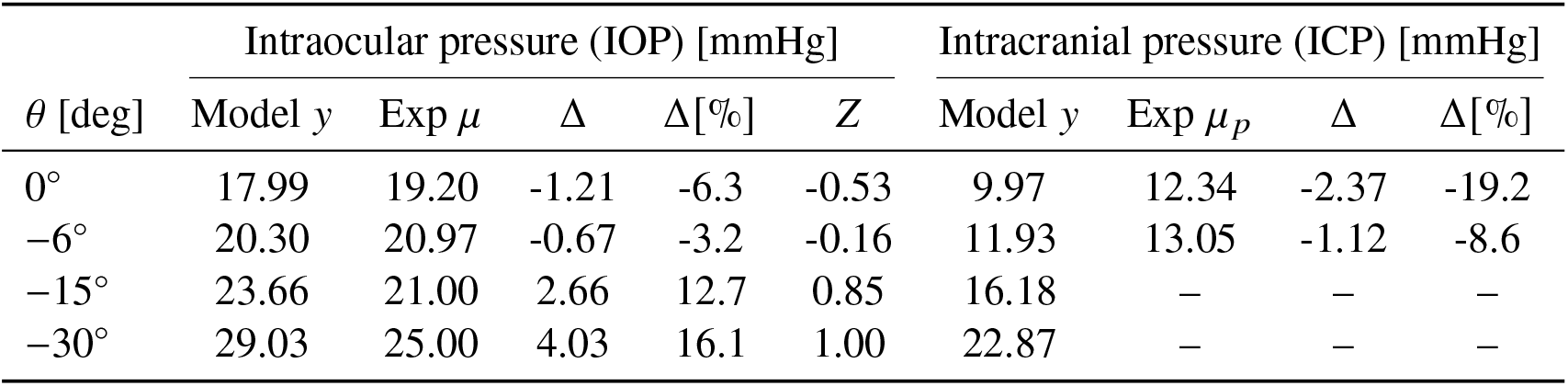
Quantitative comparison between HEAD model predictions and experimental benchmarks for intraocular pressure (IOP) and intracranial pressure (ICP) across postural tilt angles. Δ and Δ % represent the absolute and relative errors, respectively, while *Z* represents the standardized residual for IOP.

| $\theta$ [deg] | Intraocular pressure (IOP) [mmHg] | | | | | Intracranial pressure (ICP) [mmHg] | | | |
| --- | --- | --- | --- | --- | --- | --- | --- | --- | --- |
| | Model $y$ | Exp $\mu$ | $\Delta$ | $\Delta[\%]$ | $Z$ | Model $y$ | Exp $\mu_p$ | $\Delta$ | $\Delta[\%]$ |
| $0^\circ$ | 17.99 | 19.20 | -1.21 | -6.3 | -0.53 | 9.97 | 12.34 | -2.37 | -19.2 |
| $-6^\circ$ | 20.30 | 20.97 | -0.67 | -3.2 | -0.16 | 11.93 | 13.05 | -1.12 | -8.6 |
| $-15^\circ$ | 23.66 | 21.00 | 2.66 | 12.7 | 0.85 | 16.18 | — | — | — |
| $-30^\circ$ | 29.03 | 25.00 | 4.03 | 16.1 | 1.00 | 22.87 | — | — | — |

**Figure 5:**
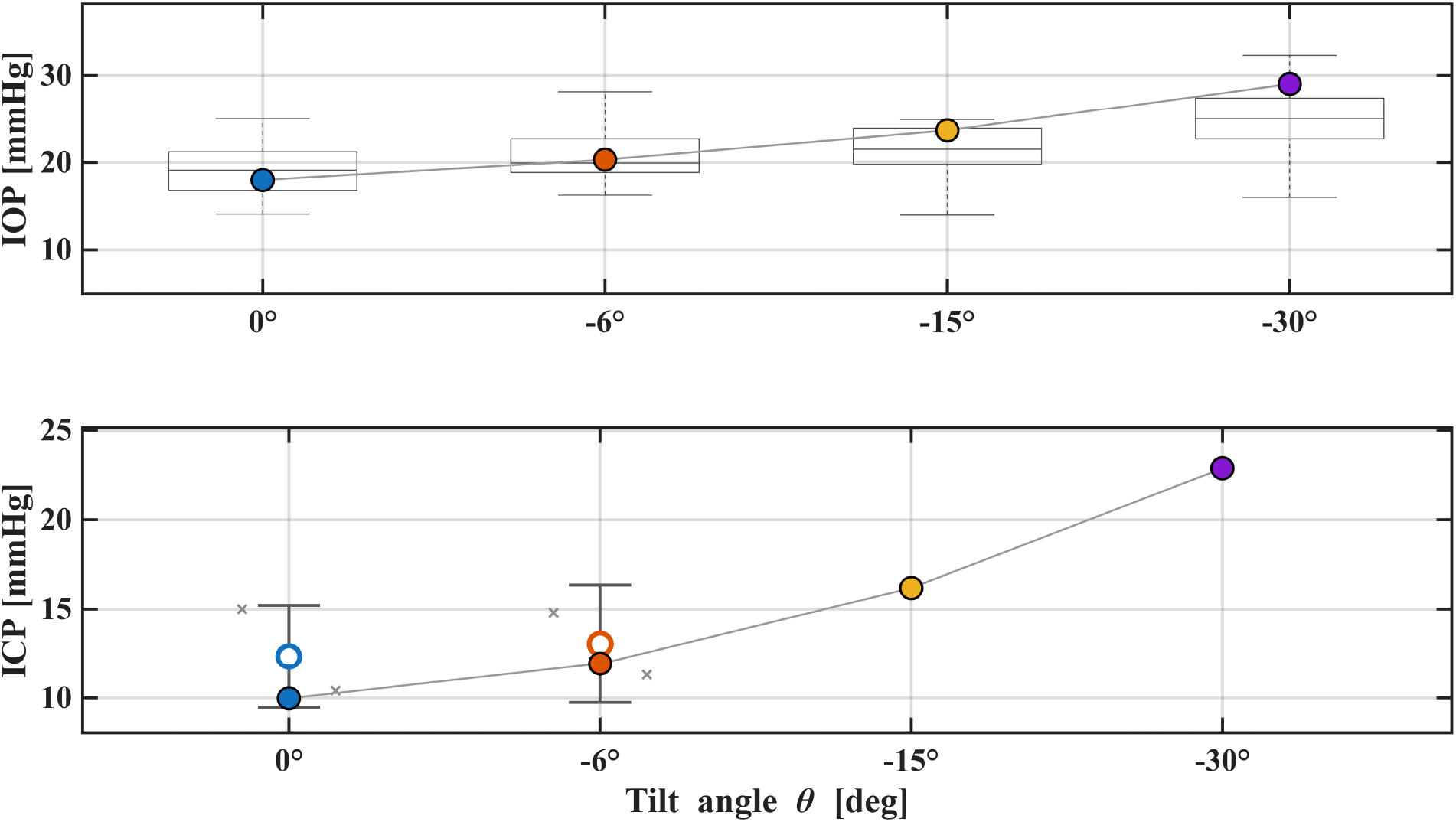
Model predictions (cycle-averaged steady-state values) are compared against experimental benchmarks at the four investigated tilt angles (*θ* = 0°, − 6°, − 15°, − 30°). On top, IOP: boxplots report subject-level measurements at each posture to preserve the experimental distribution, with the corresponding model values overlaid. On the bottom, ICP: study-level means are pooled angle-wise via sample-size weighting and shown with their ± 1 standard deviation (± 1*σ*); model predictions are superimposed at the corresponding angles.

For IOP, the model reproduces the expected monotonic increase with head-down tilt, from 17.99 mmHg at 0° to 29.03 mmHg at −30°. Across the four postures, the overall agreement is RMSE = 2.52 mmHg and nRMSE = 11.7%. The prediction lies within ±1*σ* of the experimental mean at all four angles, and within the 95% CI at 3 out of 4 postures (the largest mismatches occur at −15° and −30°, where the model moderately overestimates IOP by ∼ 12.7% and ∼ 16.1%, respectively).

For ICP, pooled benchmarks were available at 0° and −6°. The model captures the expected increase from supine to head-down conditions. However, it underestimates the pooled means at both postures, yielding an error of Δ = −1.12 mmHg (−8.6%) at −6° and an overall RMSE = 1.85 mmHg (nRMSE = 14.6%) over the two available angles. Importantly, heterogeneity among the aggregated ICP studies is substantial at 0° and −6° (high *I*^2^), supporting the inclusion of standard deviations and heterogeneity metrics alongside point-wise errors when interpreting ICP validation. Notably, when comparing the model specifically against the Laurie et al. (2017) dataset at −6° HDT (a standard bed-rest protocol angle), the agreement improves significantly, yielding an excellent match with RMSE = 0.63 mmHg and nRMSE = 5.6%.

### 3.2 Posture-driven changes in ocular hemodynamics

Figure 6 and Table 2 summarize the predicted pulsatile ocular blood flow rates in the retinal, choroidal, and ciliary beds, together with the total ocular inflow, across the four investigated tilt angles (*θ* = 0°, −6°, −15°, −30°).

**Table 2:** Cycle-averaged ocular flow rates and posture-dependent changes (Fig. 6). 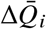 denotes the percent change relative to supine (*θ* = 0°). Pulsatility index values (PI) are reported at supine and at *θ* = −30°, together with the percent change ΔPI at *θ* = −30° vs. supine.

| Quantity | Units | $0^\circ$ | $-6^\circ$ | $-15^\circ$ | $-30^\circ$ |
| --- | --- | --- | --- | --- | --- |
| $\bar{Q}_{\text{ret}}$ | $\text{mL min}^{-1}$ | 0.04128 | 0.04266 | 0.04409 | 0.04683 |
| $\Delta\bar{Q}_{\text{ret}}$ | % (vs $0^\circ$) | - | +3.35 | +6.81 | +13.45 |
| $\bar{Q}_{\text{ch}}$ | $\text{mL min}^{-1}$ | 0.39237 | 0.40077 | 0.40192 | 0.40342 |
| $\Delta\bar{Q}_{\text{ch}}$ | % (vs $0^\circ$) | - | +2.14 | +2.43 | +2.82 |
| $\bar{Q}_{\text{cil}}$ | $\text{mL min}^{-1}$ | 0.13744 | 0.14094 | 0.14415 | 0.15065 |
| $\Delta\bar{Q}_{\text{cil}}$ | % (vs $0^\circ$) | - | +2.55 | +4.88 | +9.61 |
| $\bar{Q}_{\text{tot}}$ | $\text{mL min}^{-1}$ | 0.57109 | 0.58438 | 0.59015 | 0.60090 |
| $\Delta\bar{Q}_{\text{tot}}$ | % (vs $0^\circ$) | - | +2.33 | +3.34 | +5.22 |
| | | PI at $0^\circ$ | | PI at $-30^\circ$ ($\Delta\text{PI}$) | |
| Retinal | — | 0.398 |  | 0.350 (−12.1%) |  |
| Choroidal | — | 0.377 |  | 0.367 (−2.4%) |  |
| Ciliary | — | 0.391 |  | 0.348 (−11.0%) |  |
| Total | — | 0.382 |  | 0.361 (−5.4%) |  |

**Figure 6:**
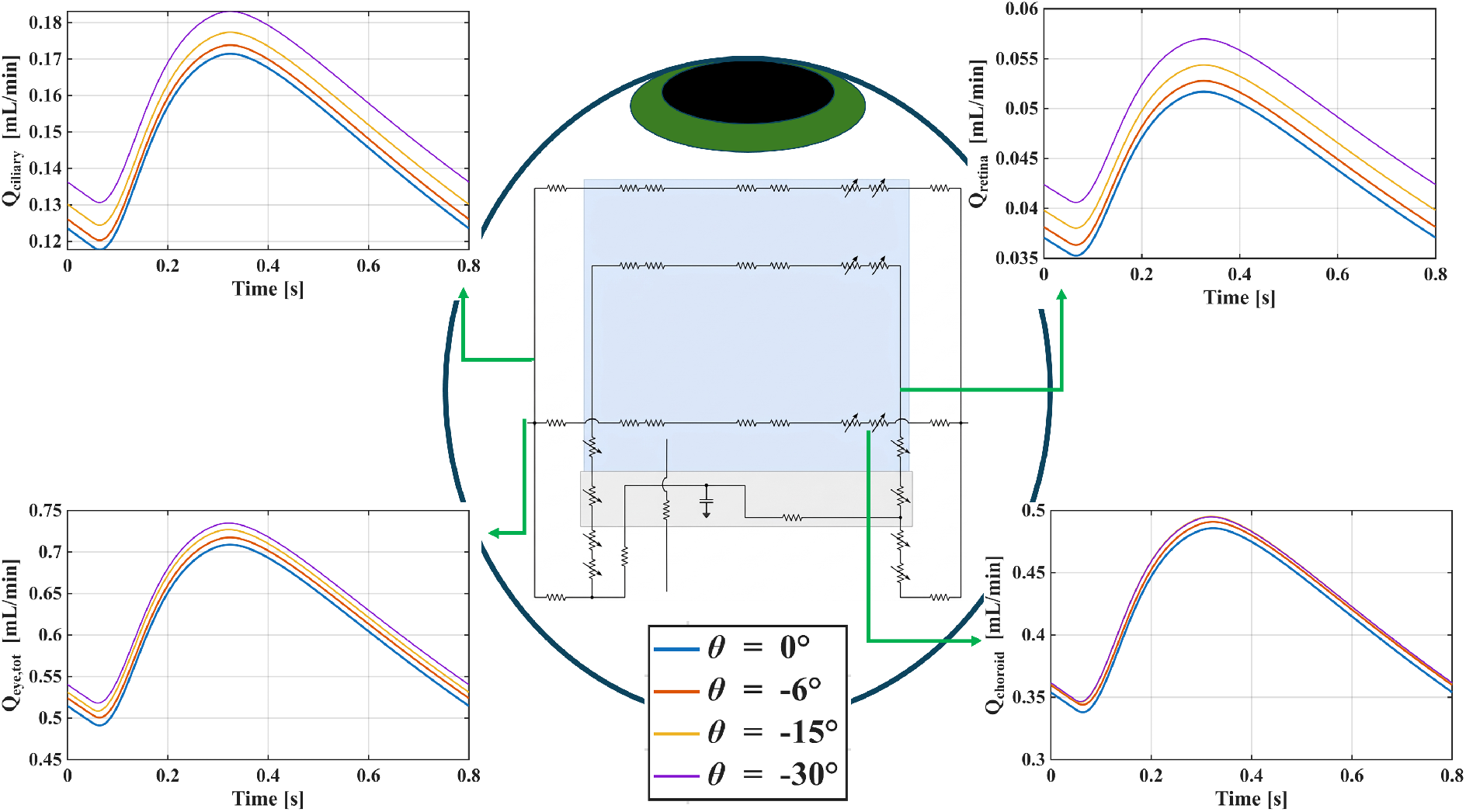
The four panels report the steady-state waveforms over the last cardiac cycle of the ciliary *Q*_ciliary_, retinal *Q*_retina_, choroidal *Q*_choroid_, and total ocular *Q*_eye,tot_ flow rates across the investigated postures (*θ* = 0°, −6°, −15°, −30°).

Starting from a supine baseline of 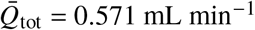, the cycle-averaged total inflow progressively increased under head-down tilt. It reached 0.584, 0.590, and 0.601 mL min^−1^ at *θ* = −6°, −15°, and −30°, respectively, which corresponds to increments of +2.33%, +3.34%, and +5.22% relative to the supine position (Table 2).

Bed-specific responses differed in magnitude. From *θ* = 0° to −30°, retinal flow exhibited the largest relative increase (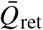 from 0.01413 to 0.00468 mL min^−1^, +13.45%), followed by the ciliary circulation (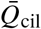 from 0.137 to 0.151 mL min^−1^, +9.6%), whereas choroidal flow showed a smaller relative change (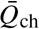 from 0.392 to 0.403 mL min^−1^, +2.8%). All three vascular beds increased monotonically with increasing head-down tilt angle.

Despite the larger relative change in the retinal bed, the absolute increase in total inflow from *θ* = 0° to −30°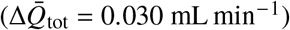 was dominated by the ciliary and choroidal circulations, accounting for approximately 44% and 37% of the total increment, respectively, with the retinal contribution limited to ∼ 19%.

Regarding waveform morphology, the pulsatile flow traces (Fig. 6) primarily exhibited an upward shift of the baseline level with increasing head-down tilt, while maintaining nearly posture-invariant peak-to-trough amplitudes: the absolute pulse amplitude (maximum minus minimum flow) remained within ±2.5% of the supine value across all beds and tilt conditions. This upward shift with approximately constant pulsatile amplitude led to a reduction in relative pulsatility under head-down tilt. At *θ* = −30°, the pulsatility index 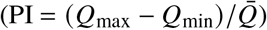 decreased by 5.4% for the total inflow relative to supine, with the largest reductions occurring in the retinal (−12.1%, from PI = 0.398 to 0.350) and ciliary (−11.0%, from PI = 0.391 to 0.348) beds, while the choroidal pulsatility index exhibited only a minor decrease (−2.4%, from PI = 0.377 to 0.367) (Table 2).

### 3.3 ONSAS compartment: CSF exchange and translaminar metrics

Figure 7 and Table 3 summarize the CSF-mediated eye–brain coupling enabled by modelling the ONSAS as a distinct compliant compartment. Across postures, mean ICP increased markedly from 9.97 mmHg at *θ* = 0° to 22.87 mmHg at *θ* = −30°, while ONSAS pressure rose from 6.28 to 20.98 mmHg. Importantly, *P*_ONSAS_ did not coincide with ICP, with a residual compartmentalization of Δ(ICP − *P*_ONSAS_) = 1.88– 3.69 mmHg depending on posture (Table 3), although the relative coupling strengthened at steeper head-down tilt (i.e., *P*_ONSAS_ approached ICP at *θ* = −30°).

**Table 3:**
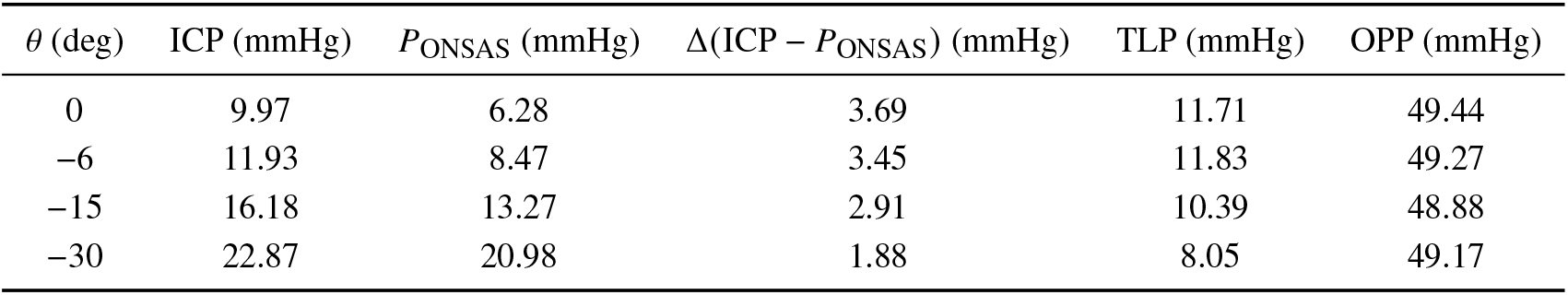
ONSAS compartment pressures and translaminar metrics across posture (cycle-mean values).

| $\theta$ (deg) | ICP (mmHg) | $P_{\text{ONSAS}}$ (mmHg) | $\Delta(\text{ICP} - P_{\text{ONSAS}})$ (mmHg) | TLP (mmHg) | OPP (mmHg) |
| --- | --- | --- | --- | --- | --- |
| 0 | 9.97 | 6.28 | 3.69 | 11.71 | 49.44 |
| −6 | 11.93 | 8.47 | 3.45 | 11.83 | 49.27 |
| −15 | 16.18 | 13.27 | 2.91 | 10.39 | 48.88 |
| −30 | 22.87 | 20.98 | 1.88 | 8.05 | 49.17 |

**Figure 7:**
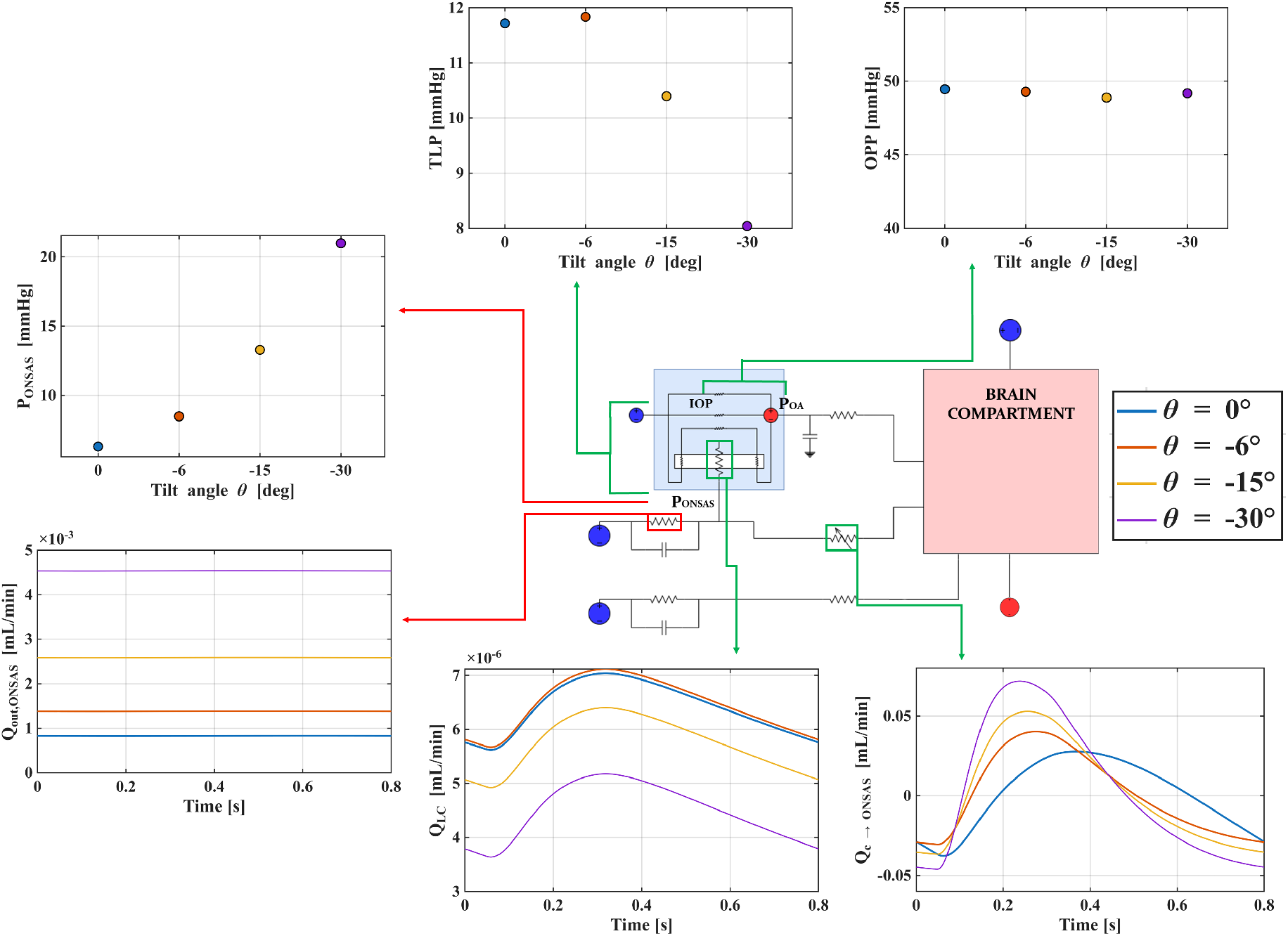
Top panels report cycle-mean pressures and derived quantities across tilt (brain compartment vs ONSAS compartment), highlighting the posture-dependent rise of ICP and *P*_ONSAS_ and their residual decoupling (ICP ≠ *P*_ONSAS_), together with the resulting translaminar pressure TLP = IOP − *P*_ONSAS_ and ocular perfusion pressure. Bottom panels show representative steady-state waveforms over the last cardiac cycle for *P*_ONSAS_ and the CSF exchange flow rates through the ONSAS pathways, including the cranial-to-ONSAS exchange, the lamina-cribrosa exchange *Q*_LC_, and the ONSAS outflow *Q*_out,ONSAS_.

This pressure redistribution translated into a reduction of translaminar loading. The translaminar pressure (TLP = IOP − *P*_ONSAS_) was 11.71 mmHg in the supine position, increased slightly to 11.83 mmHg at *θ* = −6°, and then decreased to 8.05 mmHg at *θ* = −30° (Table 3). Notably, within the simulated range, TLP remained positive, i.e. no inversion was observed. OPP remained nearly unchanged across supine and head-down conditions (48.88–49.44 mmHg), indicating that posture-driven arterial-pressure changes were largely compensated by the concurrent IOP elevation.

Beyond static levels, the model predicts posture-dependent CSF exchanges through the ONSAS pathway (Fig. 7). The ONSAS outflow *Q*_out,ONSAS_ exhibited the strongest monotonic trend, increasing from 8.27 × 10^−4^ at *θ* = 0° to 4.53 × 10^−3^ mL/min at *θ* = −30°. In contrast, the lamina cribrosa exchange flow *Q*_LC_ remained approximately two to three orders of magnitude smaller (∼ 4–6 × 10^−6^ mL/min) with only a modest decrease at steeper head-down tilt. The cranial-ONSAS exchange flow *Q*_*c*→ONSAS_ retained its sign across postures, while its cycle-mean magnitude dropped from the supine baseline (1.35 × 10^−2^ mL/min) and plateaued during head-down tilt (∼ 5.6–6.5 × 10^−3^ mL/min), concomitant with an increased pulsatile component (peak-to-peak from 6.54 × 10^−2^ to 1.18 × 10^−1^ mL/min). All posture-resolved waveforms and the full set of CSF exchange variables (including flow rates and pulsatility metrics) are provided in the Supplementary Material.

## 4 Discussion

The present study introduces the HEAD model, a unified lumped-parameter framework integrating cerebrovascular, ocular hemodynamic, and CSF dynamics under gravitational loading. The model builds upon our previous computational framework [16] by introducing a fully coupled eye-brain-CSF representation and explicitly accounting for gravitational effects and body orientation. The framework further integrates and consistently couples several physiological models previously proposed in the literature [25, 18, 28, 15, 21], bringing them together within a single comprehensive eye-brain-CSF model. This unified approach enables the simultaneous analysis of ICP, IOP, ocular blood flow redistribution, ONSAS pressure, and translaminar metrics across different body orientations. Across the four simulated tilt conditions, the model reproduces the experimentally observed increase in IOP during head-down tilt and captures the main ICP trends reported in the literature. Importantly, the coupled formulation predicts a posture-dependent pressure difference between the ICP and ONSAS, as well as heterogeneous ocular hemodynamic responses across the retinal, choroidal, and ciliary circulations. To the best of our knowledge, the proposed model is the first to explicitly account for the interaction between IOP, ICP, and ONSAS under gravitational loading, providing new insights into how eye–brain–CSF coupling shapes the translaminar mechanical environment under gravitational stress and contributes to the mechanisms underlying SANS and related ocular pathologies.

### 4.1 Validation of IOP and ICP predictions

The model captures the monotonic IOP increase from 17.99 mmHg (0° supine) to 29.03 mmHg (−30° HDT), consistent with the Petersen et al. [15] tilt dataset and the Fois et al. [14] HDT protocol, with all predictions falling within ±1*σ* of the experimental mean, yielding an overall RMSE of 2.52 mmHg and an nRMSE of 11.7%. A moderate overestimation emerges at steep HDT (+12.7% at −15°; +16.1% at −30°). For ICP, when comparing the model against the Laurie et al. [34] dataset at −6° HDT (the standard bed-rest protocol angle), it achieves an excellent match with an RMSE of 0.63 mmHg (nRMSE = 5.6%). Across the two available pooled benchmarks (0° and −6°), the overall RMSE is 1.85 mmHg. This prediction error is well within the inter-subject physiological variability (∼ 2.7 mmHg) reported by Linden et al. [35] across various tilt positions.

Importantly, the ICP pooled benchmarks at 0° and −6° exhibit substantial between-study variability (*I*^2^ = 96.7% and 83.3%). Such dispersion likely reflects not only physiological variability across cohorts but also differences in measurement techniques, calibration procedures, and experimental protocols, all of which can introduce systematic and random errors in ICP estimation. Consequently, a relevant portion of the observed variance cannot be attributed solely to sampling uncertainty, and the model-data discrepancies at these angles should be interpreted in light of these methodological limitations.

Overall, the combined IOP and ICP validation across the four tilt conditions provides a comprehensive assessment for a coupled cerebro-ocular framework. The level of agreement achieved is considered adequate to support the subsequent analysis of ocular hemodynamics and CSF dynamics presented below.

### 4.2 Ocular hemodynamic response to head-down tilt

A key advantage of the fully coupled architecture is the ability to resolve bed-specific ocular flow responses that emerge from the interaction between upstream cerebrovascular autoregulation, posture-dependent IOP, and local vascular properties; quantities that cannot be accessed by single-subsystem models.

Starting from the supine baseline, total ocular blood flow exhibits modest but progressive increases under HDT (+5.2% from 0° to −30°) as autoregulatory mechanisms partially compensate the continued rise in driving pressure.

The predicted differences among the vascular beds reflect the complex interplay between their specific autoregulatory capacity and their local biomechanical exposure to posture-induced IOP changes. Retinal flow shows the largest relative increase (+13.5%), the ciliary circulation shows a moderate increase (+9.6%), and the choroidal circulation shows the smallest change (+2.8%).

The retinal circulation possesses robust myogenic and metabolic autoregulation without autonomic innervation [26, 36], whereas the choroidal circulation, richly innervated but with limited intrinsic autoregulation [36, 37], responds largely passively to perfusion pressure changes. The model predicts that the choroidal circulation responds more passively than the retinal one, with changes that more closely track the posture-dependent variation in perfusion pressure. This trend is qualitatively consistent with laser Doppler flowmetry measurements by Longo et al. [38] and Kaeser et al.[39].

The progressive decrease in pulsatility index under HDT (retinal −12.1%, ciliary −11.0%, choroidal −2.4% at −30° vs. supine) is qualitatively consistent with experimental observations of decreased pulsatile ocular blood flow during HDT [40] and with color Doppler studies showing decreased resistive indices in the supine position [41]. In the model, the mechanism is the upward shift of mean flow with approximately constant absolute pulse amplitude, leading to a reduction in relative pulsatility. This differential pulsatility response across beds represents a novel model prediction that could be tested experimentally using bed-resolved imaging techniques such as OCT-angiography during controlled tilt protocols.

### 4.3 ONSAS compartmentalization and translaminar pressure

The treatment of the ONSAS as a distinct bilateral compartment, the central modeling innovation of this work, yields a pressure that consistently differs from ICP, with a gap Δ(ICP − *P*_*ONSAS*_) ranging from 3.69 mmHg in supine to 1.89 mmHg at −30° HDT. This compartmentalization is supported by the anatomical evidence of Killer et al. [42], by biochemical concentration gradients between lumbar and ONSAS CSF [43], and by direct pressure measurements in canine models showing *P*_*ONSAS*_ consistently lower than ICP [44]. The predicted magnitude falls within the range estimated by Holmlund et al. [45] (0–4.8 mmHg depending on sheath distensibility). The gap narrows with increasing HDT, reflecting the progressive distension of the optic nerve sheath under elevated CSF pressure, which reduces the Darcy resistance and facilitates pressure equilibration.

The resulting TLP profile is slightly non-monotonic across the simulated postures: TLP is 11.71 mmHg in the supine position, exhibits a minor peak at −6° (11.83 mmHg), and then decreases to 8.05 mmHg at −30°. This behavior arises because IOP and *P*_*ONSAS*_ respond differently to posture: at mild HDT, IOP rises faster than *P*_*ONSAS*_ due to the episcleral venous pressure increase and the compartmentalization delay; at steep HDT, *P*_*ONSAS*_ equilibrates more fully with ICP and its rapid rise outpaces the IOP increase, compressing TLP. The prediction that TLP remains positive at all angles is consistent with the in vivo measurements by Eklund et al. [46]. While Eklund et al. carefully adjusted their lumbar CSF pressure recordings to account for hydrostatic gradients and estimate ICP at the level of the lamina cribrosa, they inherently assumed a perfect pressure transmission to the retrolaminar space (i.e., ICP = *P*_*ONSAS*_). According to the present model, even after adjusting for hydrostatic differences, equating cranial ICP with *P*_*ONSAS*_ systematically overestimates the retrolaminar pressure due to the flow resistance and compartmentalization of the optic nerve sheath. This distinction highlights the added value of the compartmentalized ONSAS representation: models and clinical protocols that neglect the physiological pressure drop between the intracranial compartment and the ONSAS may misestimate the actual mechanical loading at the optic nerve head.

The absence of absolute TLP inversion in our predictions refines the hypothesis of Berdahl et al. [47]; it suggests that SANS-related optic disc changes do not require an actual pressure reversal, but can arise solely from the chronic, sustained TLP reduction caused by the loss of diurnal posture cycling in microgravity.

On Earth, the estimated 24-hour average TLP (∼ 17.3 mmHg [46]) substantially exceeds the sustained value of ∼ 8–12 mmHg predicted under continuous HDT, suggesting that the loss of diurnal posture cycling in microgravity may chronically flatten the translaminar pressure environment. Finally, the model predicts that the translaminar CSF exchange *Q*_LC_ is two to three orders of magnitude smaller than the cranio–ONSAS bulk flow *Q*_*c*→ONSAS_. This is consistent with emerging evidence [48] that the lamina cribrosa acts as a high-resistance barrier to extracellular bulk flow, restricting transport to a highly regulated, pressure-driven pathway. This prediction is uniquely enabled by the coupled framework’s simultaneous resolution of ocular and CSF pressures on both sides of the lamina.

## 5 Limitations and future developments

The HEAD model is formulated as a lumped-parameter representation of the coupled eye-brain system. This approach provides a computationally efficient framework to study the interactions between cerebrovascular circulation, ocular hemodynamics, and CSF dynamics under varying gravitational conditions. As a consequence of this spatially averaged formulation, the model does not resolve local three-dimensional fluid–structure interactions or tissue-level mechanical heterogeneity, such as lamina cribrosa deformation or detailed stress distributions along the optic nerve sheath. Nevertheless, the framework captures the global pressure and flow dynamics governing the eye–brain system and provides physiologically consistent boundary conditions (IOP, *P*_*ONSAS*_, *P*_*LC*_, ocular blood flows) for downstream high-resolution biomechanical analyses of the optic nerve head [28]. Future work will focus on coupling the present framework with structural or poroelastic models within a multiscale modeling strategy.

A second modeling assumption concerns the global cardiovascular coupling. The current framework operates as an open system with prescribed arterial inflow and venous outflow pressures. Open-loop formulations are commonly used in lumped-parameter models of ocular and cerebrovascular physiology, as they allow subsystem interactions to be analyzed under controlled boundary conditions and maintain consistency with the original physiological models on which the present framework builds. While this architecture is appropriate for investigating acute tilt conditions, it does not capture the full spectrum of systemic cardiovascular adaptations associated with prolonged exposure to microgravity, such as fluid volume redistribution and baroreflex-mediated regulation. Future work will focus on coupling the HEAD model with closed-loop cardiovascular models to enable self-consistent simulations of long-duration microgravity conditions.

Another limitation concerns the representation of subject variability. The current baseline parameter set is calibrated for a standard healthy male subject (∼ 80 kg, 170 cm) and therefore does not explicitly account for inter-individual differences in ocular or perioptic anatomy, nor for other physiological factors that may influence susceptibility to SANS [49, 50]. Future developments will focus on incorporating subject-specific parameterization based on clinical imaging datasets and parameter estimation techniques, enabling the extension of the framework toward predictive, patient-specific simulations.

Finally, the coupled architecture of the model opens several translational perspectives not explored in the present study. The framework could be used to evaluate potential SANS countermeasures *in silico*, such as lower-body negative pressure or venoconstrictive thigh cuffs, by modifying venous or gravitational boundary conditions and predicting the resulting changes in TLP, OPP, and ocular blood-flow distribution. Moreover, the explicit representation of the ONSAS as a dynamically distinct CSF compartment may also prove relevant for terrestrial pathologies such as normal-tension glaucoma and idiopathic intracranial hypertension, where the translaminar pressure balance plays a central pathogenic role [7].

Despite the limitations discussed above, the HEAD model captures the essential pressure driven interactions between the cerebrovascular, ocular, and CSF compartments under varying gravitational conditions. The model integrates cerebrovascular dynamics, multi-territory ocular hemodynamics, and posture-dependent ONSAS compartmentalization within a unified lumped-parameter formulation.

## 6 Conclusions

This work presents the HEAD model, a unified lumped-parameter framework that integrates cerebrovascular dynamics with autoregulation, multi-territory ocular hemodynamics, and compartmentalized craniospinal-ONSAS cerebrospinal fluid circulation into a single system of nonlinear ODEs. By dynamically linking the subsystems governing the eye-brain pressure environment, the model enables a self-consistent analysis of ICP, IOP, ocular blood flow, and CSF dynamics under gravitational variations, providing new insights into the mechanisms underlying SANS and related ocular pathologies.

Model simulations reproduce the main posture-dependent trends in ICP and IOP observed experimentally and reveal heterogeneous ocular hemodynamic responses across retinal, choroidal, and ciliary circulations. Importantly, representing the ONSAS as a CSF compartment dynamically distinct from the intracranial space leads to a posture-dependent decoupling between ICP and perioptic pressure, shaping the translaminar pressure environment at the optic nerve head.

These results highlight the importance of eye-brain-CSF coupling in determining the mechanical environment across the lamina cribrosa under gravitational stress. In this context, the HEAD model provides a quantitative framework to investigate how gravitationally driven fluid shifts reshape the coupled eye–brain pressure environment, offering new mechanistic insights into the development of SANS, while also providing physiologically grounded boundary conditions for future structural or transport models of the optic nerve head. Its modular architecture further enables future extensions toward long-duration microgravity simulations, coupling with systemic cardiovascular models, and subject-specific investigations of pressure-related ocular pathologies.

## Data availability statement

All generated raw simulation data and the experimental datasets used for model validation are available on Figshare (DOI: 10.6084/m9.figshare.31779646).

## Author contributions

**Conceptualization**: Michele Nigro, Eduardo Soudah. **Methodology**: Michele Nigro, Eduardo Soudah. **Formal Analysis**: Michele Nigro. **Software**: Michele Nigro. **Validation**: Michele Nigro. **Funding acquisition**: Eduardo Soudah. **Writing-original draft**: Michele Nigro, Andrea Montanino, Eduardo Soudah. **Writing-review & editing**: Michele Nigro, Andrea Montanino, Eduardo Soudah.

